# Fecal bacterial communities of the platypus (*Ornithorhynchus anatinus*) reflect captivity status – implications for conservation and management

**DOI:** 10.1101/2023.12.04.570006

**Authors:** Ashley M. Dungan, Jessica L. Thomas

## Abstract

The duck-billed platypus (Ornithorhynchus *anatinus*) is currently listed as ‘Near-Threatened’ under the International Union for Conservation of Nature (IUCN) Red List based on observed population declines and local extinctions. A key part of the conservation strategy for this species is its captive maintenance; however, captive animals often undergo significant changes in their gut microbiome. The study of the gut microbiome in threatened wildlife species has enormous potential to improve conservation efforts and gain insights into host-microbe coevolution. Here, for the first time, we characterize the gut microbiome of wild platypus via fecal samples using high-throughput sequencing of the bacterial 16S rRNA gene and identify microbial biomarkers of captivity in this species. At the phylum level, Firmicutes (50.4%) predominated among all platypuses, followed by Proteobacteria (28.7%), Fusobacteria (13.4%), and Bacteroidota (6.9%), with twenty-one ‘core’ bacteria identified. Captive individuals did not differ in their microbial α-diversity compared to wild platypus but had significantly different community composition (β-diversity) and exhibited higher abundances of *Enterococcus*, which are potential pathogenic bacteria. Four taxa were identified as biomarkers of wild platypus, including *Rickettsiella, Epulopiscium, Clostridium*, and Cetobacterium. This contrast in gut microbiome composition between wild and captive platypus is an essential insight for guiding conservation management as the rewilding of captive animal microbiomes is a new and emerging tool to improve captive animal health, maximize captive breeding efforts, and give reintroduced or translocated animals the best chance of survival.

## 1 Introduction

The platypus (*Ornithorhynchus anatinus*), is a semiaquatic, egg-laying mammal endemic to eastern Australia, including Tasmania. Platypuses have many Aboriginal names including Mallangong, Tambreet, Gaya-dari, Boonaburra, and Lare-re-lar [1,2]. Indigenous Australians were the first to describe the biology of the platypus and developed a deep biocultural and ecological knowledge of this animal, which was largely overlooked by early naturalists. A Dreamtime story of the platypus from the Darling (Barka) River [1] begins with a young duck who ventures down the creek far from her tribe. She is abducted by Biggoon, a large water rat who took the duck as his wife. The duck eventually escaped and returned to her tribe, where she laid two eggs that hatched as platypuses. They had soft fur instead of feathers, four webbed feet instead of two, and spurs on their hind legs, like Biggoon’s spear. The duck and her two different children were banished by her tribe, choosing to live far away in the mountains where she could hide from her tribe and Biggoon.

The platypus is the sole living representative of its family (Ornithorhynchidae) and genus (*Ornithorhynchus*), though several related species appear in the fossil record. This species was listed as ‘Near-Threatened’ under the International Union for Conservation of Nature (IUCN) Red List in 2016 based on observed population declines and local extinctions [3], though significant uncertainty exists about its current distribution and abundance [4]. Large-scale, intense conservation efforts, such as animal captivity and species translocation, have been put forward to accelerate ecosystem recovery and reduce species declines. Captivity provides a suitable opportunity to care for threatened species [5] and can facilitate reintroduction efforts. However, dietary changes, habitat homogenization, stress, intra- and interspecific interactions, and antibiotic usage can all contribute to microbial dysbiosis, increased prevalence of disease, and reduced reproductive success in captivity [6-8]. This highlights the critical need to develop and identify effective methods that can improve animal wellbeing in captivity, with manipulation of the host microbiome being an emerging, possibly powerful tool for species conservation [8].

In mammals, the transfer of bacteria from mother to offspring is crucial for the establishment of a healthy gut microbiota, immune system development, protection against pathogens, and metabolic health [9]. Once established, gut microbiota influence immunity, energy uptake from different diet types, detoxification of plant secondary compounds, protection from pathogens, chemical communication, behaviour, and stress responses of the mammalian host [10-12]. Because microbiomes can encompass a hundred-fold more genes than host genomes and respond to the environment over both daily and evolutionary time scales, microbiomes function as a phenotypically plastic buffer between the host-genotype’s effects and the environmental effects that interact to shape host phenotypes [11,13,14]. As such, there are several opportunities for transfer of bacteria from the mother to her young (vertical transmission) and from the environment (horizontal transmission).

The reproductive process of the platypus presents an opportunity for vertical transmission of bacteria from the female to the egg, as the cloaca is used to both excrete waste (feces and urine) and lay eggs [15]. In reptiles, for example, vertically transmitted bacteria from the cloaca protects the egg from fungal infections and egg failure [16]. During lactation, milk is secreted through ducts in the mammary glands and is absorbed directly through openings in the skin on the abdomen. These openings allow the milk to be drawn into the mouths of the young platypuses (known as puggles) as they lick and suck on the mother’s skin [17]. Further, females have been observed rubbing their bill over the bills of the nestlings after they had suckled [18]. These behaviours serve as other opportunities for the puggles to potentially acquire bacteria. Unlike eutherian neonates, Monotreme puggles are exposed to the external environment prior to developing a functional immune system and are thus highly vulnerable to microbial infections [19].

Understanding microbial communities and how they may shift is particularly relevant to endangered wildlife, such as the platypus, maintained in captivity as well as those reared in captivity for reintroduction to the wild [20], as it allows us to gain insights into host-microbe coevolution [21] and inform conservation efforts [8,22]. Here, for the first time we characterize the wild and captive gut microbiota of platypus, identifying bacterial biomarkers of wild animals that can be used in microbiome manipulation approaches.

## 2 Materials and Methods

### 2.1 Study animals

This study included samples collected from one adult male housed alone, a pair of adult females housed together, and an adult female housed alone. Each animal had access to approximately 8,000 L of potable tap freshwater and an artificial ‘burrow’ composed of timber boxes for resting. The water bodies were furnished with natural vegetation, logs and river rocks. Platypuses were fed *ad libitum* a diet of freshwater crayfish (*Cherax destructor and C. albidus*), mealworms (*Tenebrio molitor*), earthworms (*Oligochaeta* sp.), blackworms (*Lumbriculus variegatus*), and fly pupae (*Musca domestica*).

Wild platypuses were captured using fyke nets in Coranderrk creek, Victoria (DELWP permit 10009424). Animals were immediately released after microchipping and physical assessment.

### 2.2 Sample collection and processing

Due to the non-invasive nature of fecal collection, feces have been extensively used to study gut microbial composition of mammals during captivity [23-25]. Though there are differences in microbial community composition between gastrointestinal and fecal samples [26], the use of feces as a proxy for the gut microbial community has revealed important and biologically meaningful findings in both humans and animals [27,28] and is the only non-invasive method available for sample collection from wild animals. Sample collections from wild-captured and captive platypus were made under permits from The University of Melbourne Animal Ethics Committee (Permit no. 21001) and DELWP permit 10009424.

Captive fecal samples (n=17) were collected from the four resident platypuses at the Healesville Sanctuary (HS) in Victoria, Australia from December 2021 to April 2022. Samples were collected by scooping fecal material that had been deposited on the bottom of the water tank with a sterile FLOQSwab (Copan Diagnostics, Murrieta, CA, USA) in the morning during routine cleaning and transferring the swab tip to a sterile 1.5 mL tube. As most tanks have a single resident platypus, we were able to track the identity of the source animal. Wild-captured platypus fecal samples (n=5) were collected with FlOQSwabs opportunistically from annual platypus surveys in Coranderrk Creek during handling for a physical exam as per [18]. Food (mealworms, earthworms, and bloodworms; n=3 each) samples were collected by placing whole organisms in 1.5 mL tubes using sterile forceps. Enclosure water samples (n=8) were collected from each platypus tank by sampling 50 mL with a sterile 50-mL polypropylene tube. All tubes (1.5 and 50 mL) were stored at -20 °C until processing. The sample size for both captive and wild platypus samples was constrained by the small size of the captive platypus population at HS and by the opportunistic nature of the wild sampling as to not unnecessarily disturb native populations. The majority of feces are deposited in the water by the platypuses. Occasionally, feces are deposited when the animal is close to the bottom of the tank which leaves a visible sample.

### 2.3 DNA extraction and library prep

Fecal (n=22), food (n=9), and enclosure water (n=8) samples used in this project were brought to The University of Melbourne from the Healesville Sanctuary on ice. Upon arrival, enclosure water samples were filtered through a 0.22 um membrane; the membrane was collected and transferred to a sterile 2 mL tube for DNA extractions. DNA was extracted from each sample using FastDNA SPIN Kit (MP Biomedicals, Australia) for soil following the manufacturer instructions. Extraction blanks (n=3) were included as controls. Extracted DNA and controls were amplified by PCR in triplicate using primers with sequencing adapters (underlined) targeting the V4 region of the bacteria 16S rRNA gene: 515F (511 - <u>GTGACCTATGAACTCAGGAGTC</u>GTGCCAGCMGCCGCGGTAA - 3’ [29]) and 806R (511 - <u>CTGAGACTTGCACATCGCAGC</u>GGACTACHVGGGTWTCTAAT-3⍰ [30]), with three no template PCR negatives included. These primers were chosen to correspond with the Earth’s microbiome protocol and to be comparable with other native Australian animal microbiome studies [31-33]. Libraries were prepared for sequencing following Dungan, et al. [34]. Briefly, triplicate PCRs were comprised of 1 μL template DNA, 7.5 μL of 2x MyTaq HS Mix polymerase (Bioline), 0.45 μL of 10 μM forward and reverse primers, and nuclease-free water to 15 μL. Thermal cyclers were set to 1 cycle × 95°C for 3 min; 18 cycles × 95°C for 15 s, 55°C for 30s, and 72°C for 30s; 1 cycle × 72°C for 7 min; hold at 4°C. Triplicate PCR products were then pooled; successful DNA extraction was confirmed by agarose gel electrophoresis.

A volume of 20 μL of each PCR product pool was purified by size-selection using Nucleomag NGS Clean-up and Size Select beads (Scientifix, Australia). The purified DNA was resuspended in 40 μL of nuclease-free water. Indexing PCRs were created by combining 10 μL of purified DNA with 10 μL 2x MyTaq HS Mix polymerase (Bioline) and 1 μl (5 μM) of forward and reverse indexing primers. Thermal cyclers were set to: 1 cycle × 95°C for 3 min; 24 cycles × 95°C for 15 s, 60°C for 30 s, and 72°C for 30 s; 1 cycle × 72°C for 7 min; hold at 4°C. For a subset of randomly chosen samples, product size was confirmed by agarose gel electrophoresis. A sequencing library was created by pooling 5 μL from each reaction and performing a final bead clean-up. The library was checked for quality and quantity (2200 TapeStation, Agilent Technologies, Australia) to guide pool normalization, then sequenced on a single Illumina MiSeq run using v3 (2 × 300 bp) reagents at the Walter and Eliza Hall Institute, Melbourne, Australia.

### 2.4 Metabarcoding data analysis

Raw demultiplexed 16S rRNA gene sequences were imported into QIIME2 v2022.8 [35] where primers and overhang adapters were removed (using cutadapt v2.6; [36]). The data were then filtered, denoised using the pseudo-pooling method, and chimera checked [using DADA2 v1.22.0; 37] to generate amplicon sequence variants (ASVs). Taxonomy for each ASV was assigned against a SILVA database (version 138) trained with a naïve Bayes classifier against the same V4 region targeted for sequencing [38]. A phylogenetic tree was produced in QIIME2 by aligning ASVs using the PyNAST method [39] with mid-point rooting.

### 2.5 Statistical analysis

All data were analyzed in R [v4.3.1; 40]. For metabarcoding data, ASV, taxonomy, metadata and phylogenetic tree files were imported into R and combined into a phyloseq object [41]. Contaminant ASVs were identified and removed sequentially from the dataset according to their abundance in the PCR negative controls and extraction blanks relative to the samples using the prevalence method in the R package decontam with p = 0.1 [42].

α-diversity metrics (observed ASVs, Shannon’s and inverse Simpson’s indices) were calculated from a rarefied dataset using the ‘estimate_richness’ function in the R package phyloseq [41]. These were then analyzed using linear mixed effects models using the R package nlme [43]. Where appropriate, post hoc comparisons were performed using Tukey’s honestly significant difference test in the R package emmeans [44]. Box plots were created with ggplot2 [45], plotrix [46], and gridExtra [47]. Summary data was produced using the function ‘summarySE’ from the package Rmisc [48].

β-diversity was evaluated using a Bray-Curtis dissimilarity matrix. The difference in community compositions among groups (wild versus captive) were calculated using ‘adonis2’ (a modified version of a permutational multivariate analysis of variance (PERMANOVA)) in the vegan package in R [49] with 9999 permutations. This method tests the hypothesis that the bacterial communities from different groups are significantly different from each other in terms of their composition. PERMANOVA uses a distance matrix as input and is often more robust to non-normality and heteroscedasticity than traditional ANOVA. To account for unequal samples sizes, the ‘strata’ argument was included; this helps to ensure that the PERMANOVA analysis appropriately considers the sampling imbalance.

Where the assumption of homogeneity of dispersion was not met, a permutation test for multivariate dispersion to account for unequal dispersion between groups was included. Where appropriate, Holm corrected pairwise comparisons were computed using the package pairwiseAdonis [50]. The results of the PERMANOVA were cross validated with and visualized using principal coordinates analysis (PCoA) to gain a more robust understanding of the underlying patterns.

Where the PERMANOVA test revealed that the community composition was significantly different (p < 0.05), linear discriminant analysis effect size (LEfSe) in the microbiomeMarker package [51] was used to identify bacteria taxa that were more dominant in each group than other layers based on a p < 0.05 from the Wilcox test and LDA score >4.0. The cladogram illustrated the taxonomic relationship of the associated with captivity status. Barplots visualizing overall microbiota composition were made using ggplot2 [45] by agglomerating taxa at the genus or ASV level based on relative abundances. Venn diagrams were created using packages eulerr [52] and microbiome [53]; upset plots were generated using the packages UpSetR [54] and MicrobiotaProcess [55].

## 3 Results

### 3.1 Dataset overview

Sequencing resulted in 2.7 M reads across all platypus fecal samples (n=22), water samples (n=8), platypus feed (n=9), extraction blanks (n=3), and PCR negative control samples (n=3). After filtering, denoising, and removal of chimeras, 1.6 M reads remained, and samples averaged 36,535 reads. ASVs with fewer than 10 reads were removed from the dataset as they are not biologically feasible. Decontam identified eight putative contaminant ASVs from PCR amplification (0.58% of total reads) and fifteen putative contaminant ASVs from DNA extraction (1.73% of total reads) (Table S1), which were removed from the dataset. One wild platypus sample had fewer than 100 reads and was removed from analysis. After all filtering steps, there were 1661 ASVs across the remaining samples (n = 38) for downstream analysis in R.

### 3.2 Abundant and core microbiota

The largest taxonomic groups in the platypus gut microbiome were an unknown *Lactobacillales* (17.1%), *Fusobacteriaceae* (13.3%), *Enterobacteriaceae* (10.7%), *Peptostreptococcaceae* (9.3%), *Enterococcaceae* (7.3%), and *Clostridiaceae* (6.7%), members of the phyla Fusobacteriota (13.4%), Firmicutes (50.4%), and Proteobacteria (28.7%) (Fig. 1A; Fig. S1). Twenty-one core ASVs that averaged ≥0.1% abundance in ≥ 70% of platypus (both captive and wild) samples were identified (Table S2). These ASVs are largely members of the dominant families, but also include two representatives each from *Vagococcaceae, Streptococcaceae*, and *Morganellaceae*.

**Figure 1.**
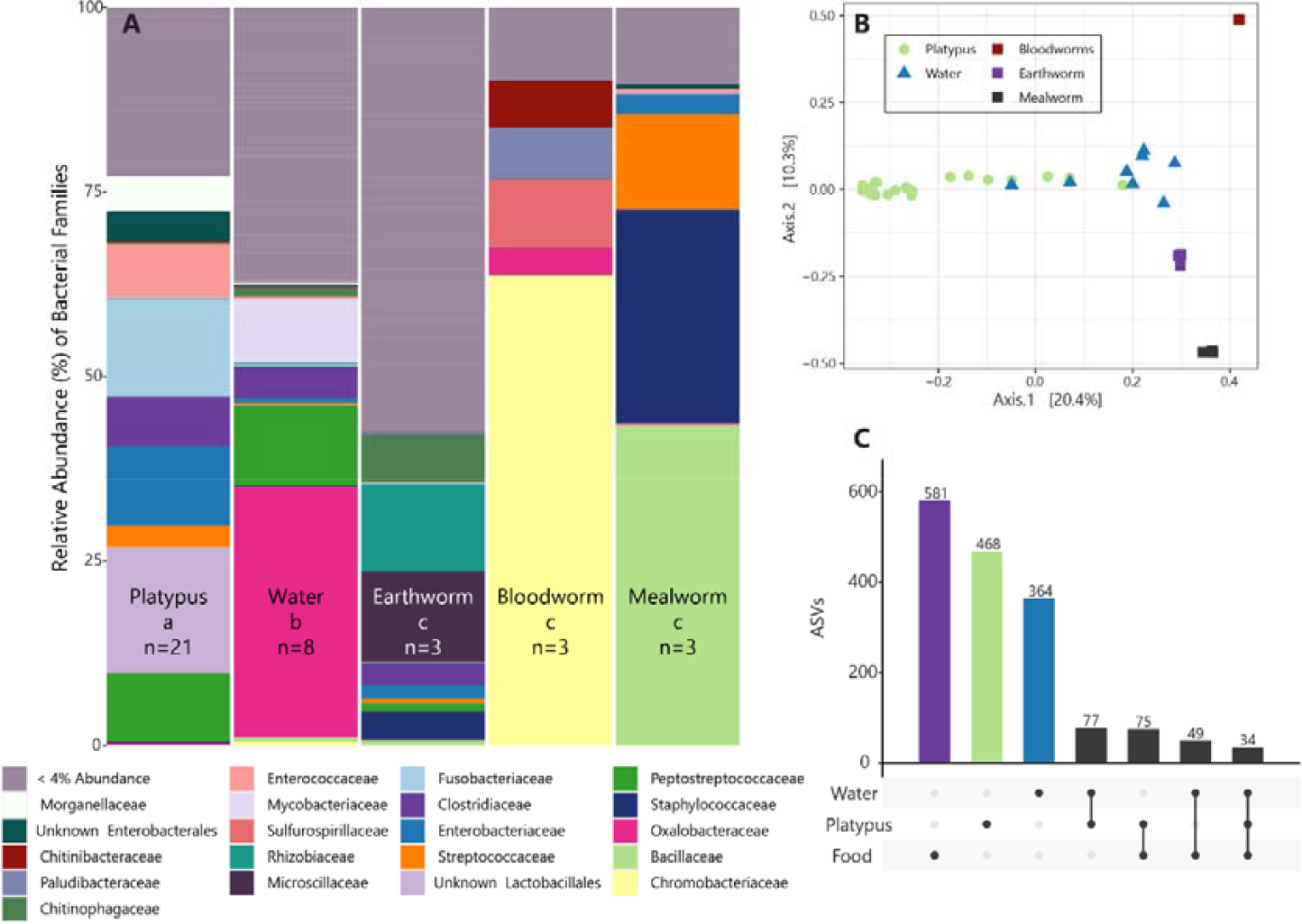
**(A)** Relative abundance of bacterial families in the platypus fecal microbiome (captive and wild), enclosure water, and three types of feed (earth, blood, and mealworms). Families whose relative abundance was less than 4% in all sample types were pooled into the “< 4% Abundance” category. Different small letters under each sample type indicate significant differences in Tukey’s honest significant difference (HSD) post hoc tests for β-diversity as measured by PERMANOVA (pairwise stats are presented in Table S3). **(B)** Principal coordinate analysis (PCoA) for all sample types using a Bray-Curtis dissimilarity matrix with platypus (circle), enclosure water (triangle), and food sources (square). **(C)** UpsetR plot illustrating the number of shared and unique ASVs for enclosure water (blue), platypus (wild and captive; green), and feed (earth-, blood-, and mealworms; purple) samples.

Platypus-associated microbial communities (β-diversity) were significantly different from the surrounding aquatic environment and food sources (F_(4,37)_=5.82, p=0.0001; Fig. 1); these different environments explain >40% of the variation in the microbiome composition (R^2^=0.41345). Pairwise comparisons showed that the platypus microbiome is distinct from all food sources and enclosure water, which were also significantly different from one another (Table S3; Fig. 1). Of the dominant microbial taxa, Peptostreptococcaceae and Clostridiaceae were found in the enclosure water at 10.8% and 4.3% relatively abundance, respectively. Clostridiaceae (3.0% - Earthworm) and Enterobacteriaceae (2.7% - Mealworm; 1.8% - Earthworm) were also found in food sources, although at lower relative abundances. While there were 34 shared ASVs between platypus, enclosure water, and food sources, there were a considerable number of unique ASVs with 468, 364, and 581 distinct ASVs for platypus, water, and food samples, respectively (Fig. 1C). Samples were rarefied to 7491 reads for α-diversity metrics. There were significant differences in richness (F_(4,29)_=16.73, p<0.0001; Fig. 2A) and Shannon diversity (F_(4,29)_=8.60, p=0.0001; Fig. 2C) according to sample type (Table S4), but no significant differences in Simpson’s index of diversity.

**Figure 2.**
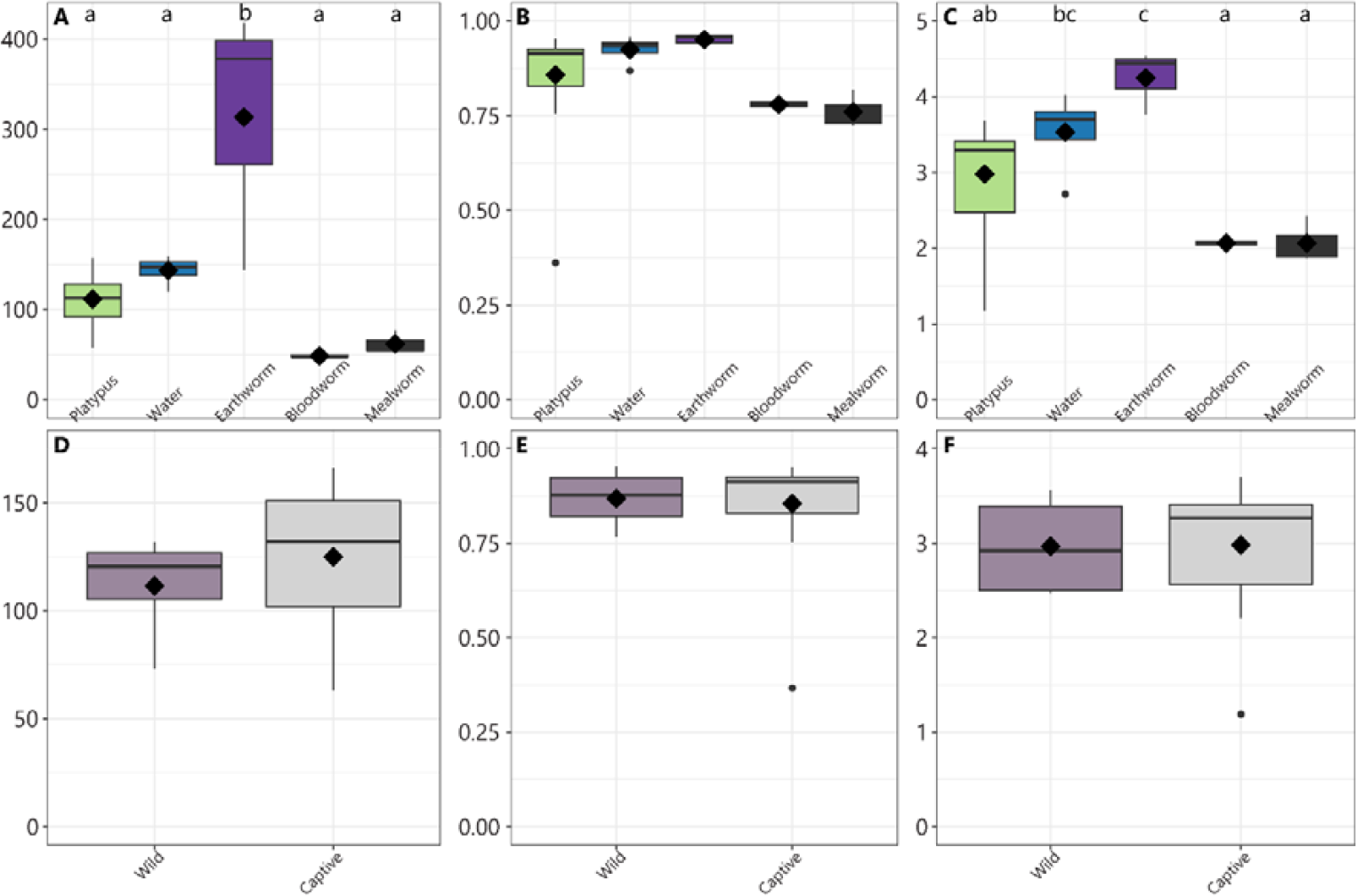
**(A-C)** α-diversity metrics for all samples by category and **(D-F)** for platypus only by captivity status, showing **(A,D)** richness, **(B,E)** Simpson’s index of diversity, and **(C,F)** Shannon’s index. Boxes cover the interquartile range (IQR) and the diamond inside the box denotes the median. Whiskers represent the lowest and highest values within 1.5 × IQR. Different small letters signify significant differences in Tukey’s honest significant difference (HSD) post hoc tests; pairwise stats are presented in Table S4.

### 3.3 Wild versus captive platypus microbiome

There were no significant differences in α-diversity metrics (rarefied to 30,503 reads) between wild and captive individuals (Fig. 2D-F); however, there was a significant difference in microbiota composition (F_(1,20)_=2.00, p=0.0347; Fig. 3). Captivity status explains approximately 10% of the total variation (R^2^=0.09536).

**Figure 3.**
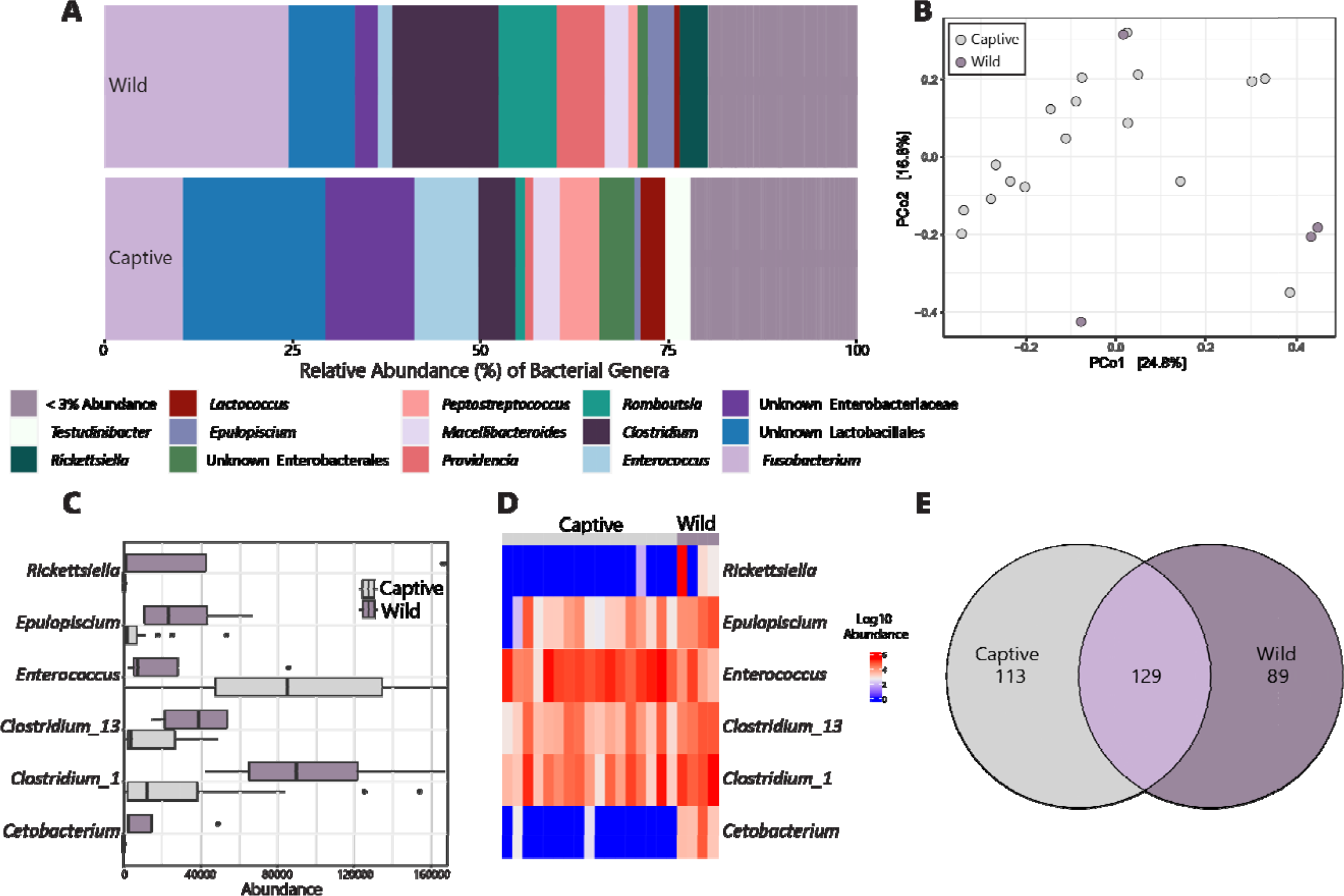
Taxonomic composition at the genus level for wild and captive platypus fecal samples **(A)**. Principal coordinate analysis (PCoA) for all sample types using a Bray-Curtis dissimilarity matrix for wild (purple) and captive (grey) platypus **(B)**. Abundance **(C)** and heatmap **(D)** of the bacterial genera that were biomarkers for wild or captive platypus determined by LEfSe based on a P < 0.05 from the Wilcox test and linear discriminant analysis score >4.0. Venn diagram illustrating the number of shared and unique ASVs for platypus (wild and captive), enclosure water, and feed (earth-, b lood-, and mealworms) where a given ASV was at least 0.1% abundant in any sample. **(E)**

Linear discriminant analysis Effect Size (LEfSe) analysis identified five genera that are biomarkers of captivity status (Fig. 3C-D; Table 1). Biomarkers for wild platypus include the taxa *Clostridium* (two variants), *Rickettsiella, Epulopiscium*, and *Cetobacterium*, while only one taxon (*Enterococcus*) was a biomarker for captive platypus.

**Table 1:**
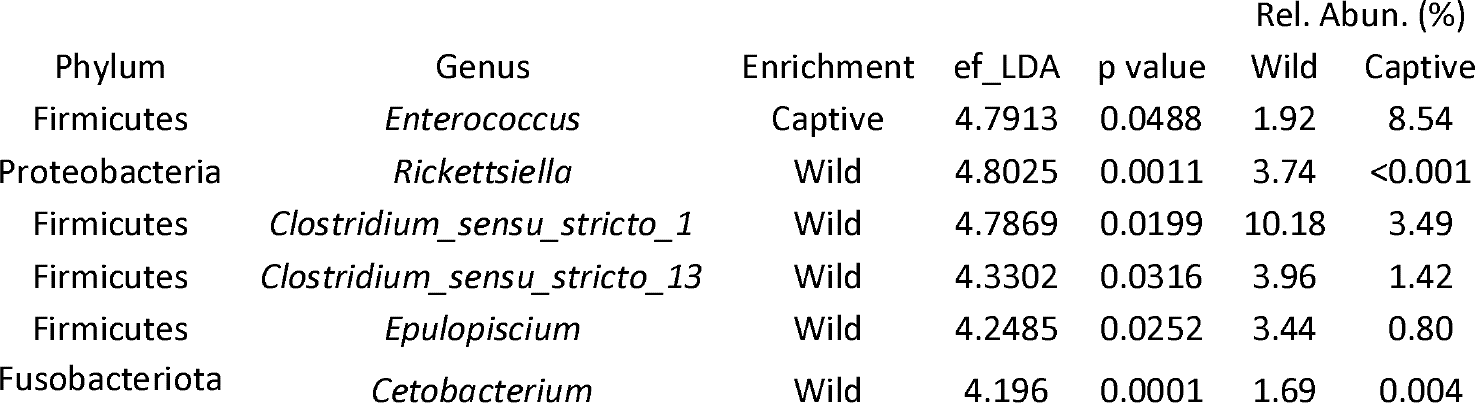
Genera identified as biomarkers for captivity status, where enrichment indicates what group the genus was a biomarker for. Linear Discriminant Analysis (LDA) is a dimensionality reduction technique used in LEfSe to identify features that have the largest differences between groups while minimizing the within-group variability. The effect size in LEfSe is calculated using LDA scores, which reflect the strength and direction of the association between a feature and a particular group (ef_LDA).

## Discussion

### Major constituents of the platypus fecal microbiome

The fecal microbiome of the platypus is dominated by an unknown *Lactobacillales, Fusobacteriaceae, Enterobacteriaceae, Enterococcaceae, Peptostreptococcaceae*, and *Clostridiaceae*, the latter two of which are also found in reasonable abundance in their habitat water. This could be an artifact of the animals defecating in their aquatic environment (horizontal transfer to water) or these taxa could be consistently present in the enclosure water and transfer to the platypus via ingestion or through direct contact with the feces. While some dominant taxa were also found in food sources (i.e., *Clostridiaceae* and *Enterobacteriaceae*), the platypus microbiome is distinct from its habitat and diet, suggesting that these microbiota are symbiotic and critical for the health of the animal. Members of the platypus microbiome are commonly found in other mammalian fecal samples, including those native to Australia such as the Tasmanian devil [56], Northern quoll [57], other semi- and fully aquatic mammals [58], and their closest living relative and fellow monotreme, the short-beaked echidna [31,59].

### Influence of diet on the platypus microbiome

It is understood that diet plays a significant role in shaping the gut/fecal microbiota in mammals [28]. In the wild, platypus feed on annelid worms, insect larvae, and bottom-dwelling crustaceans [60], and Healesville Sanctuary works to replicate this diet. Another typically opportunistic carnivore, the Tasmanian devil, has a microbiome that is also dominated by Firmicutes, Proteobacteria, Fusobacteria, and Bacteroidota [56,61]. Other striking similarities between the platypus and the Tasmanian devil gut microbiome are the relatively high proportion of Proteobacteria and Fusobacteria with lower abundances of Firmicutes and Bacteroidota compared to other mammals [62,63], which leads to a high Firmicutes to Bacteroidota ratio (F:B ratio) [56]. It has been found in humans and mice that a high F:B ratio (the “obese microbiome”) is associated with high efficiency in energy harvest from the diet [64]. Interestingly, low levels of Bacteroidota and high levels of Proteobacteria have also been observed in the gut microbiome of many other carnivorous mammals [63] besides devils and platypus, including the cheetah (*Acinonyx jubatus*) [65], spotted hyena (*Crocuta crocuta*), and polar bear (*Ursus maritimus*) [62]. These findings suggest that a high F:B ratio could be a feature of carnivorous species possibly related to the need to efficiently harvest and store energy from limited food sources [56]. In the platypus, this is in line with their feeding behaviors, whereby platypuses consume a significant proportion of their body weight each day, with daily food intake ranging from 13-19% and potentially reaching 90-100% during late lactation [60,66]. During egg laying and immediately following, female platypus may go reduced feeding for 2-3 weeks [60]; a microbiome that supports the efficient use and storage of nutrients is critical for their survival. From the wild to captive individuals, we observed an increase in the proportion of Bacteroidota (4.3% in wild vs. 7.5% in captive; Fig. S1) resulting in a shift from a 10.5:1 F:B ratio in wild samples to a 6.5:1 ratio in captive animals. Changes to the F:B ratio may reflect changes in the pattern of food intake by platypus during captivity, where sources of food are more consistent compared to wild animals.

### Captivity alters the platypus microbiome

Relative to wild populations, the generalized pattern of gut microbiomes in captivity are reduced α-diversity and a significant shift in community composition [8]. Here, we saw no changes in α-diversity, though this may have been an artefact of low replication for wild samples, with an alteration in bacterial community structure. Many conditions of captivity (antibiotic exposure, altered diet composition, homogenous environment, increased stress, and altered intraspecific interactions) likely lead to changes in the host-associated microbiome [67-70].

LEfSe determined biomarkers for wild platypus include the taxa *Clostridium* (two variants), *Rickettsiella, Epulopiscium,* and *Cetobacterium*, while only one taxon (*Enterococcus*) was a biomarker for captive platypus. *Clostridium*, or its parent family Clostridiaceae, have previously been identified as a biomarker in wild mammalian [71] and reptile [72] microbiomes, with potential roles in degrading fiber [73] and proteins [74]. In dairy cows, a high relative abundance of *Epulopiscium* in the fecal microbiota was attributed to their plant-based diet [75], as this taxa produces specialized enzymes to help digest complex polysaccharides associated with plants and algae [76]. Thus, the presence of higher abundances of *Clostridium and Epulopiscium* in wild-captured platypus could be vital for breaking down and utilizing various complex plant-derived polysaccharides, which would be an incidental constituent of their diet as they opportunistically feed on bottom-dwelling aquatic organisms. *Cetobacterium* has also been identified as a biomarker for wild Tasmanian devils [56] and waterbirds [77]. Members of this genus are common in the gastrointestinal tract of marine and aquatic mammals and have the ability to produce vitamin B_12_, which is essential for the biosynthesis of haem, a component of haemoglobin and myoglobin, but not produced by mammals [78].

Not all biomarkers for wild platypus have beneficial roles. In Australia, *Rickettsiella* is an obligate intracellular parasite and common microbiome member in ticks [79]. Ticks potentially act as reservoirs for this zoonotic bacterium, passing it on to wild platypus. This taxon is responsible for diseases in wild animals such as lymphadenopathy and spotted fever [80].

*Enterococcus*, a facultatively anaerobic bacterium that is commonly found in mammalian gut microbiota [81], was the sole taxon identified as a biomarker of captivity. In host associations, *Enterococcus* are involved in the production of vitamins, the degradation of complex carbohydrates from the diet, and for modulation of the immune system [82]. However, Enterococci are also seen as opportunistic pathogens, due to the ease of acquiring different virulence factors and resistance to various classes of antibiotics [82]. This highlights the potential for *Enterococcus* to cause infections and become resistant to antibiotics.

### Implications for conservation efforts

One of the effective strategies to conserve endangered species with small population sizes is through translocation of individuals kept in captivity into the wild. This can introduce some challenges as the captive animals are often kept under conditions that can be very different compared with their natural environments (*e.g.*, different diets), which can have profound impacts on their microbiota [31]. Through various functions in nutrition, chemical signalling, immunity, reproduction, and host health [10-12], the animal gut microbiome can increase the ability to adjust from conditions under captivity to natural conditions.

Bioaugmentation of the microbiome through probiotic therapy, fecal microbiome transplantation, or diet modification [83] is a new and emerging field in microbiome research, especially within the context of wildlife conservation [84,85]. Augmenting or manipulating the microbiome may provide numerous benefits, such as restoring dysbiotic microbiomes for improved physiological functions and animal health [86,87], ‘re-wilding’ the microbiomes of captive animals [88], and mitigating disease risk [89]. Given the non-invasive nature of fecal collections, fecal microbiome transplantation of wild platypus to their captive counterparts could be a viable option to reintroduce potentially beneficial members of *Clostridium, Epulopiscium*, and *Cetobacterium* that were virtually absent in the captive platypus microbiome. Further, the captive diet of this species could be augmented to include riparian plants, an incidental component of the platypus diet, to assist the proliferation of *Clostridium and Epulopiscium*. Another option is probiotic administration, either of these specific bacterial candidates or with other taxa to shift microbiota composition. For example, probiotic dosage of *Bacillus subtilis* and B. ceres increased the relative abundance of *Cetobacterium* in tilapia [90].

## Conclusions

Today, platypus populations continue to face increasing pressure from habitat destruction, river regulation, netting and pollution [91]. Under current climate and threats, platypus abundance is predicted to decline by 47%–66% over the next 50 years [4]. There is an urgent need to implement national conservation efforts for this unique mammal by increasing surveys, tracking trends, mitigating threats, improving management of platypus habitat in rivers, and supporting captive populations. Given the complexities associated with managing these threats, establishing healthy insurance populations ex situ through captive breeding is essential for preventing further mammal extinctions in Australia [92]. To minimize the problems arising from captivity, efforts can be taken to manipulate microbial diversity and composition to be comparable with wild populations through methods such as increasing dietary diversity, exposure to natural environmental reservoirs, or probiotics [93]. For individuals destined for reintroduction, these strategies can prime the microbiota to buffer against novel pathogens and changes in diet and improve reintroduction success. The microbiome is a critical component of animal physiology and its role in species conservation should be expanded and included in the repertoire of future management practices.

## Declarations

### Ethics approval and consent to participate

Not applicable

### Consent for publication

Not applicable

## Availability of data and material

Raw MiSeq data are available under NCBI BioProject ID PRJNA971672. QIIME2 and R code can be found at https://github.com/adungan31/platypus.

## Authors’ contributions

AMD conceived the project; AMD and JT designed the study; JT collected collected all platypus fecal samples; AMD processed samples through metabarcoding library prep; AMD analysed and interpreted the data; AMD wrote the first draft; all authors substantively revised the manuscript and approve of the submitted version.

## Acknowledgements

The platypus whose fecal material was used in this study were originally collected from the lands of the Wurundjeri People. We acknowledge their contributions to this research not only by way of these beautiful creatures, but also through their Yarns built from lived experiences, ontologies, epistemologies and axiologies. Indigenous Knowledge is the foundation of our learning and knowledge and has inspired and guided this research. We are grateful to Drs. Justin Maire and Kshitij Tandon for reviewing this manuscript.

