## supplemental material for "Fecal bacterial communities of the platypus (*Ornithorhynchus anatinus*) reflect captivity status – implications for conservation and management"

**This file includes Figure S1 and Tables S1-S4.**

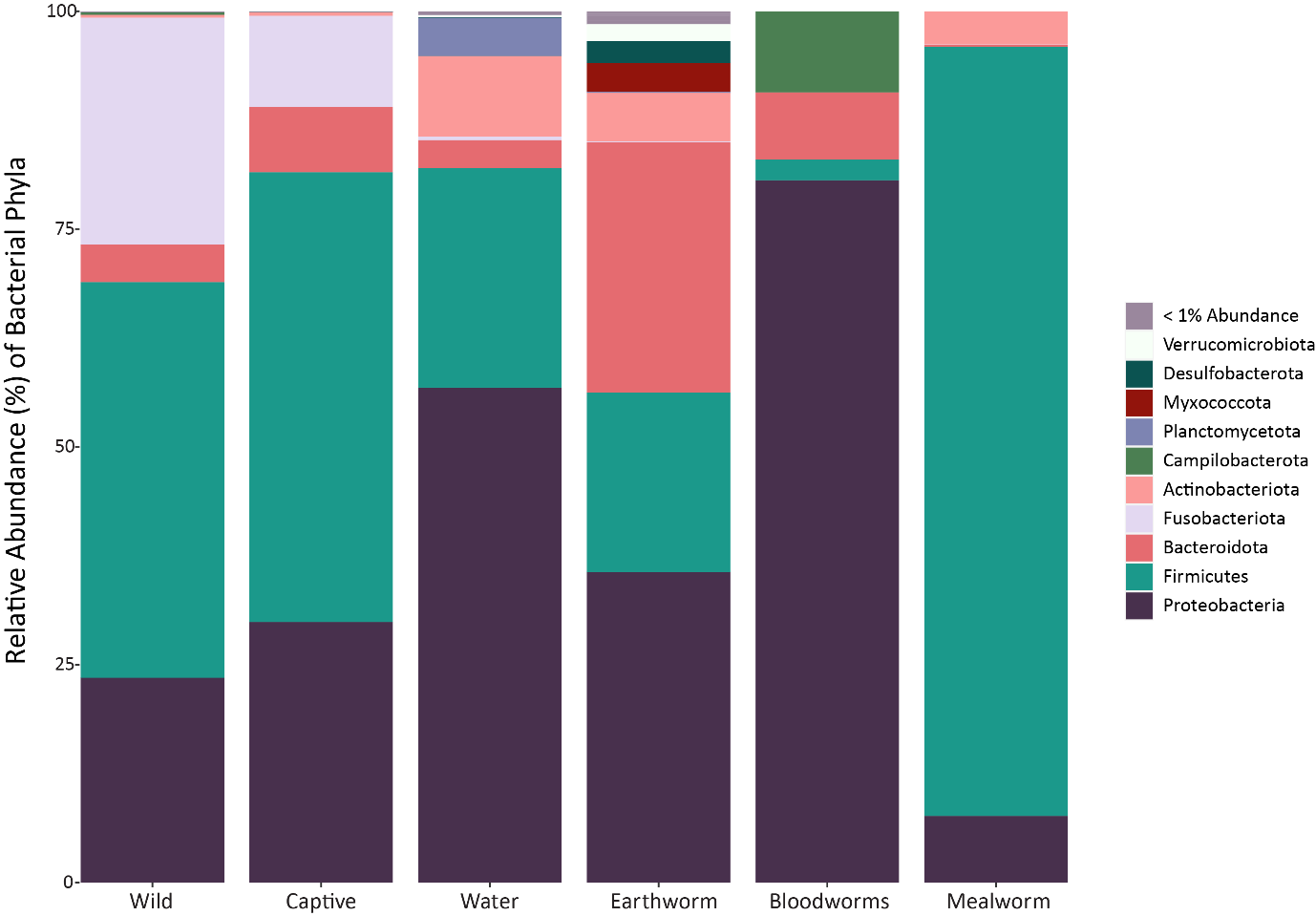

Figure S1: Relative abundance of bacterial phyla for captured and wild platypus, their enclosure water, and three types of feed (earth, blood, and mealworms). Phyla whose relative abundance was less than 1% all sample types were pooled into the “<1% Abundance” category.

Table S1: A total of 23 potential contaminants were identified from extraction blanks (n=15; 1.73% of total reads) and PCR negative controls (n=8; 0.58% of total reads) using the prevalence method in the R package decontam (Davis, et al., 2018). The relative abundance (%) of each ASV in the full dataset prior to their removal is reported in the last column.

| ASV | Source | Phylum | Class | Family | Genus | Rel. Abun. |
| --- | --- | --- | --- | --- | --- | --- |
| Contam01 | PCR Negative | Actinobacteriota | Actinobacteria | Nocardiaceae | *Rhodococcus* | 0.011 |
| Contam02 | PCR Negative | Proteobacteria | Gammaproteobacteria | Pseudomonadaceae | *Pseudomonas* | 0.006 |
| Contam03 | PCR Negative | Proteobacteria | Gammaproteobacteria | Pseudomonadaceae | *Pseudomonas* | 0.023 |
| Contam04 | PCR Negative | Proteobacteria | Gammaproteobacteria | Methylophilaceae | *Methylophilus* | 0.005 |
| Contam05 | PCR Negative | Proteobacteria | Gammaproteobacteria | Moraxellaceae | *Acinetobacter* | 0.017 |
| Contam06 | PCR Negative | Proteobacteria | Gammaproteobacteria | Halomonadaceae | *Halomonas* | 0.348 |
| Contam07 | PCR Negative | Proteobacteria | Gammaproteobacteria | Halomonadaceae | *Halomonas* | 0.160 |
| Contam08 | PCR Negative | Proteobacteria | Gammaproteobacteria | Halomonadaceae | *Halomonas* | 0.013 |
| Contam09 | Extraction Blank | Bacteroidota | Bacteroidia | Crocinitomicaceae | *Fluviicola* | 0.353 |
| Contam10 | Extraction Blank | Bacteroidota | Bacteroidia | Flavobacteriaceae | *Flavobacterium* | 0.037 |
| Contam11 | Extraction Blank | Bacteroidota | Bacteroidia | Paludibacteraceae | *Paludibacter* | 0.400 |
| Contam12 | Extraction Blank | Actinobacteriota | Actinobacteria | Micrococcaceae | *Kocuria* | 0.002 |
| Contam13 | Extraction Blank | Firmicutes | Bacilli | Staphylococcaceae | *Staphylococcus* | 0.023 |
| Contam14 | Extraction Blank | Proteobacteria | Alphaproteobacteria | Caulobacteraceae | *Brevundimonas* | 0.002 |
| Contam15 | Extraction Blank | Proteobacteria | Alphaproteobacteria | Caulobacteraceae | *Brevundimonas* | 0.005 |
| Contam16 | Extraction Blank | Proteobacteria | Alphaproteobacteria | Rhizobiaceae | *Shinella* | 0.001 |
| Contam17 | Extraction Blank | Proteobacteria | Alphaproteobacteria | Sphingomonadaceae | Unknown Sphingomonadaceae | 0.005 |
| Contam18 | Extraction Blank | Proteobacteria | Alphaproteobacteria | Sphingomonadaceae | *Sphingomonas* | 0.024 |
| Contam19 | Extraction Blank | Proteobacteria | Gammaproteobacteria | Pseudomonadaceae | *Pseudomonas* | 0.001 |
| Contam20 | Extraction Blank | Proteobacteria | Gammaproteobacteria | Oxalobacteraceae | *Janthinobacterium* | 0.759 |
| Contam21 | Extraction Blank | Proteobacteria | Gammaproteobacteria | Vibrionaceae | *Vibrio* | 0.067 |
| Contam22 | Extraction Blank | Proteobacteria | Gammaproteobacteria | Moraxellaceae | *Acinetobacter* | 0.020 |
| Contam23 | Extraction Blank | Proteobacteria | Gammaproteobacteria | Moraxellaceae | *Acinetobacter* | 0.028 |
|  |  |  |  |  | **Total** | **2.309** |

Table S2: Core taxa of the platypus faecal microbiome, identified as ASVs present in at least 70% of the samples and averaging > 0.01% relative abundance. The average relative abundance of each ASV is listed separately for captive and wild platypus populations.

|  | Classification | | | | | Rel. Abun. | |
| --- | --- | --- | --- | --- | --- | --- | --- |
| ASV | Phylum | Class | Order | Family | Genus | Captive | Wild |
| ASV001 | Fusobacteriota | Fusobacteriia | Fusobacteriales | Fusobacteriaceae | *Fusobacterium* | 6.69 | 7.01 |
| ASV002 | Firmicutes | Clostridia | Clostridiales | Clostridiaceae | *Clostridium_sensu_stricto_1* | 1.07 | 1.34 |
| ASV003 | Firmicutes | Clostridia | Clostridiales | Clostridiaceae | *Clostridium_sensu_stricto_13* | 1.42 | 1.03 |
| ASV004 | Firmicutes | Clostridia | Peptostreptococcales-Tissierellales | Peptostreptococcaceae | *Peptostreptococcus* | 5.22 | 1.20 |
| ASV005 | Firmicutes | Clostridia | Peptostreptococcales-Tissierellales | Peptostreptococcaceae | *Romboutsia* | 0.30 | 5.41 |
| ASV006 | Firmicutes | Clostridia | Peptostreptococcales-Tissierellales | Peptostreptococcaceae | *Terrisporobacter* | 0.67 | 0.56 |
| ASV007 | Firmicutes | Clostridia | Peptostreptococcales-Tissierellales | Peptostreptococcaceae | *Terrisporobacter* | 0.43 | 0.27 |
| ASV008 | Firmicutes | Bacilli | Lactobacillales | Enterococcaceae | *Enterococcus* | 3.02 | 1.09 |
| ASV009 | Firmicutes | Bacilli | Lactobacillales | Enterococcaceae | *Enterococcus* | 0.98 | 0.23 |
| ASV010 | Firmicutes | Bacilli | Lactobacillales | Vagococcaceae | *Vagococcus* | 0.36 | 0.18 |
| ASV011 | Firmicutes | Bacilli | Lactobacillales | Vagococcaceae | *Vagococcus* | 0.99 | 0.29 |
| ASV012 | Firmicutes | Bacilli | Lactobacillales | Streptococcaceae | *Lactococcus* | 0.60 | 0.13 |
| ASV013 | Firmicutes | Bacilli | Lactobacillales | Streptococcaceae | *Lactococcus* | 2.72 | 0.62 |
| ASV014 | Firmicutes | Bacilli | Lactobacillales | Unknown Lactobacillales | Unknown Lactobacillales | 18.97 | 8.84 |
| ASV015 | Firmicutes | Bacilli | Lactobacillales | Enterococcaceae | *Enterococcus* | 0.71 | 0.05 |
| ASV016 | Firmicutes | Bacilli | Lactobacillales | Enterococcaceae | *Enterococcus* | 3.24 | 0.32 |
| ASV017 | Proteobacteria | Gammaproteobacteria | Enterobacterales | Morganellaceae | *Morganella* | 1.43 | 2.91 |
| ASV018 | Proteobacteria | Gammaproteobacteria | Enterobacterales | Enterobacteriaceae | Unknown Enterobacteriaceae | 3.03 | 0.57 |
| ASV019 | Proteobacteria | Gammaproteobacteria | Enterobacterales | Enterobacteriaceae | Unknown Enterobacteriaceae | 5.11 | 1.79 |
| ASV020 | Proteobacteria | Gammaproteobacteria | Enterobacterales | Enterobacteriaceae | Unknown Enterobacteriaceae | 2.32 | 0.03 |
| ASV021 | Proteobacteria | Gammaproteobacteria | Enterobacterales | Morganellaceae | *Proteus* | 0.60 | 1.32 |

Table S3: β-diversity pairwise comparisons by sample source, which were completed using a Bray-Curtis distance matrix and the package ‘pairwise.adonis’. p values have been corrected based on a Holm adjustment method. Tukey HSD letters were assigned to each genotype based on these statistics and added to barplots in Fig. 1A.

| pairs | SumsOfSqs | F.Model | R2 | adjusted p value |
| --- | --- | --- | --- | --- |
| Platypus vs Earthworm | 1.544 | 6.145 | 0.218 | 0.004 |
| Platypus vs Mealworm | 1.830 | 7.693 | 0.259 | 0.005 |
| Platypus vs Bloodworms | 1.960 | 8.293 | 0.274 | 0.004 |
| Platypus vs Water | 1.440 | 4.671 | 0.147 | 0.002 |
| Earthworm vs Mealworm | 1.142 | 12.560 | 0.758 | 0.300 |
| Earthworm vs Bloodworms | 1.334 | 16.146 | 0.801 | 0.300 |
| Earthworm vs Water | 1.034 | 2.697 | 0.231 | 0.037 |
| Mealworm vs Bloodworms | 1.479 | 161.872 | 0.976 | 0.300 |
| Mealworm vs Water | 1.272 | 3.626 | 0.287 | 0.037 |
| Bloodworms vs Water | 1.260 | 3.632 | 0.288 | 0.037 |

Table S4: α-diversity pairwise comparisons by sample source. p values have been corrected based on a Holm adjustment method. Tukey HSD letters were assigned to each genotype based on these statistics and added to barplots in Fig. 2A-C.

| α-diversity metric | contrast | estimate | SE | df | t.ratio | p.value |
| --- | --- | --- | --- | --- | --- | --- |
| **ObsASVs** | **Bloodworms - Earthworm** | **-264.7** | **37.4** | **29** | **-7.068** | **<.0001** |
| ObsASVs | Bloodworms - Mealworm | -13.3 | 37.4 | 29 | -0.356 | 0.9964 |
| ObsASVs | Bloodworms - Platypus | -63.2 | 28.3 | 29 | -2.232 | 0.1965 |
| ObsASVs | Bloodworms - Water | -94.8 | 35 | 29 | -2.708 | 0.0772 |
| **ObsASVs** | **Earthworm - Mealworm** | **251.3** | **37.4** | **29** | **6.712** | **<.0001** |
| **ObsASVs** | **Earthworm - Platypus** | **201.5** | **28.3** | **29** | **7.118** | **<.0001** |
| **ObsASVs** | **Earthworm - Water** | **169.8** | **35** | **29** | **4.849** | **0.0003** |
| ObsASVs | Mealworm - Platypus | -49.9 | 28.3 | 29 | -1.761 | 0.4144 |
| ObsASVs | Mealworm - Water | -81.5 | 35 | 29 | -2.327 | 0.1654 |
| ObsASVs | Platypus - Water | -31.6 | 25 | 29 | -1.265 | 0.7141 |
| **Shannon** | **Bloodworms - Earthworm** | **-2.181** | **0.463** | **29** | **-4.713** | **0.0005** |
| Shannon | Bloodworms - Mealworm | 0.001 | 0.463 | 29 | 0.002 | 1 |
| Shannon | Bloodworms - Platypus | -0.912 | 0.35 | 29 | -2.606 | 0.0955 |
| **Shannon** | **Bloodworms - Water** | **-1.467** | **0.433** | **29** | **-3.388** | **0.0161** |
| **Shannon** | **Earthworm - Mealworm** | **2.182** | **0.463** | **29** | **4.715** | **0.0005** |
| **Shannon** | **Earthworm - Platypus** | **1.269** | **0.35** | **29** | **3.629** | **0.0089** |
| Shannon | Earthworm - Water | 0.714 | 0.433 | 29 | 1.65 | 0.479 |
| Shannon | Mealworm - Platypus | -0.913 | 0.35 | 29 | -2.609 | 0.095 |
| **Shannon** | **Mealworm - Water** | **-1.468** | **0.433** | **29** | **-3.39** | **0.016** |
| Shannon | Platypus - Water | -0.555 | 0.309 | 29 | -1.795 | 0.3956 |
